# Expansion Spatial Transcriptomics

**DOI:** 10.1101/2022.10.25.513696

**Authors:** Yuhang Fan, Žaneta Andrusivová, Yunming Wu, Chew Chai, Ludvig Larsson, Mengxiao He, Liqun Luo, Joakim Lundeberg, Bo Wang

## Abstract

Capture array-based spatial transcriptomics methods have been widely used to resolve gene expression in diverse tissue contexts, however, their spatial resolution is limited by array density. We present Expansion Spatial Transcriptomics (Ex-ST) to overcome this limitation by clearing and expanding tissue prior to capturing the entire transcriptome. This approach allows us to achieve near cellular resolution and higher capture efficiency of lowly-expressed genes, which we demonstrate using mouse brain samples.

## Main

The spatial distribution of transcripts is essential for understanding cell states and cellular organization in tissues. In recent years, many methods for spatial profiling of gene expression have been developed ^1,2^. Whereas imaging-based approaches including *in situ* hybridization and *in situ* sequencing require prior selection of targets, capture array methods can target the entire transcriptome ^3^. The first technique, introduced by Ståhl *et al*. in 2016, now commercialized by 10x Genomics as the Visium platform, relies on capturing polyadenylated mRNAs released from tissue sections onto barcoded array surface ^4^. Its broad applicability has been demonstrated in various systems to describe developmental processes ^5^ and profile complex diseases ^6,7^. However, its spatial resolution is limited by the density of spatially barcoded arrays. Currently, the spot size is 55 μm with a center-to-center distance of 100 μm, resulting in each spot being occupied by multiple cells.

Here we present Expansion Spatial Transcriptomics (Ex-ST), an approach that allows us to overcome the resolution limit by first expanding samples embedded in a polyelectrolyte matrix before capturing mRNAs on Visium slides. We applied this method on two mouse brain regions, olfactory bulb and hippocampus, chosen for their well-described anatomically distinct patterns with clear molecular signatures. Ex-ST significantly improves the resolution and mRNA capture efficiency of the Visium array, enabling us to better resolve cell types and detect rare transcripts.

We took inspiration from expansion microscopy, in which biomolecules are anchored to polyelectrolyte gel matrices that can be expanded to increase imaging resolution ^8^. In our Ex-ST protocol (**Fig. 1a**), a tissue section is embedded in a polyacrylate gel to anchor mRNAs. After all proteins are digested, the gel undergoes ~2.5 times linear expansion in 0.1× SSC buffer. The expanded gel is stained with DAPI, placed on the Visium capture array where fluorescent images are taken to register nuclei for aligning anatomy and gene expression in later steps. A key design of Ex-ST is the use of two poly-dT probes of different lengths and melting temperatures (MT), with the shorter one (MT ~39°C) anchored in the gel and the longer one (MT > 55°C) spatially barcoded on the Visium slides. This allows us to use heat (45°C for 30 min) to release mRNAs from the gel and recapture them on the array surface (**Extended Data Fig. 1**). Reverse transcription and library preparation was then performed using a modified version of the standard Visium protocol (see **Methods**). These modifications, together with tissue clearing that facilitates mRNA transport, enhances the efficiency of transcript detection.

**Fig. 1:**
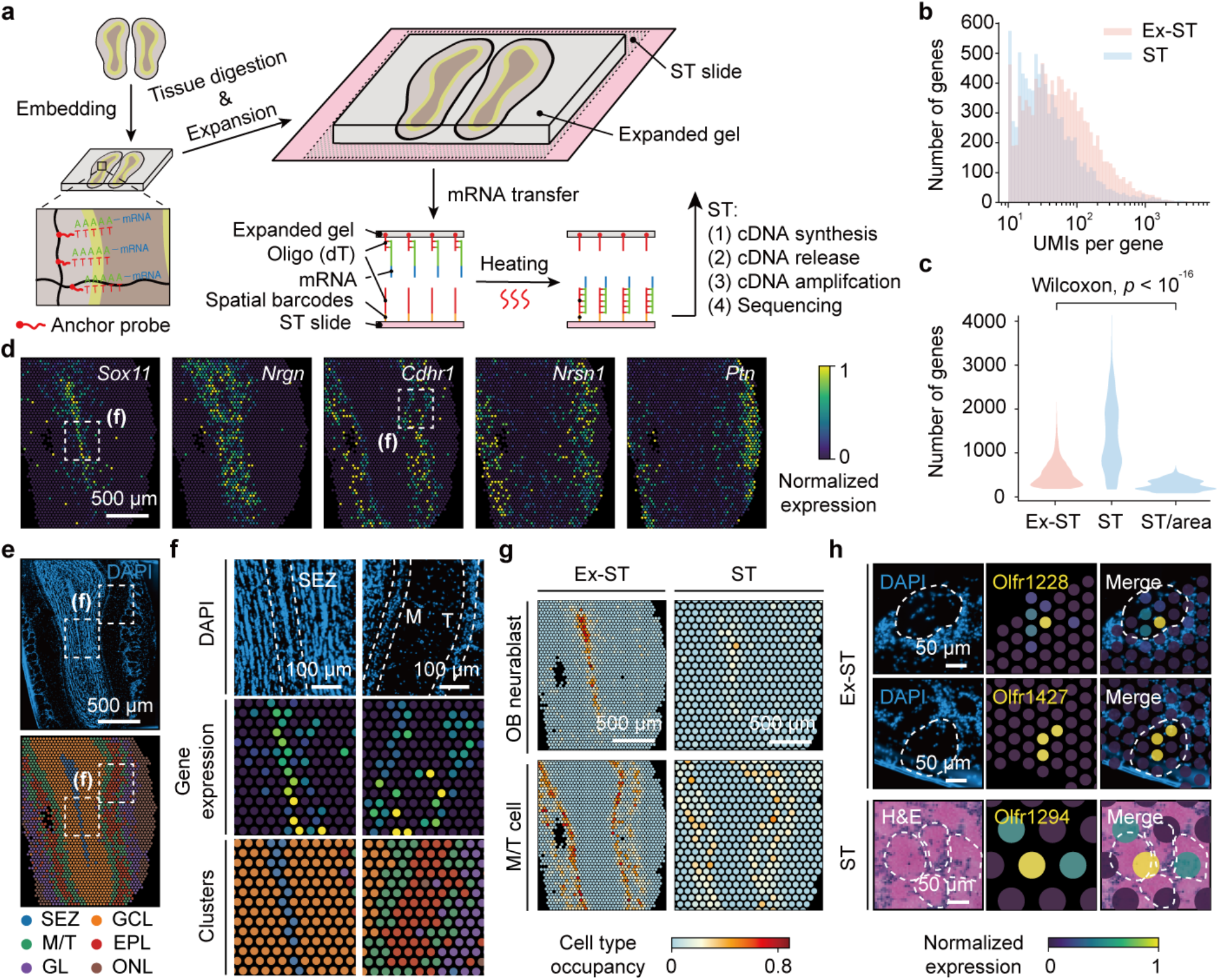
Ex-ST resolves MOB layers with high resolution. **a,** Overview of the Ex-ST workflow. **b,** Comparison of number of UMIs for all genes present in Ex-ST and standard Visium datasets, each containing two MOB sections. **c,** Comparison of number of genes per spot measured by Ex-ST, standard Visium, and standard Visium after normalizing the tissue area covering each spot. **d,** Spatially resolved expression of region-specific marker genes processed by the Ex-ST protocol. **e,** DAPI image of expanded MOB section (top) and annotated clusters on tissue coordinates (bottom): SEZ, subependymal zone; GCL, granular layer; M/T, mitral and tufted cell layer; EPL, external plexiform layer; GL, glomerular layer; ONL, olfactory nerve layer. **f,** Magnified views of SEZ and M/T regions processed by Ex-ST. Top: DAPI images with SEZ and M/T regions with dashed outlines; middle: expression of region-specific marker genes (*Sox11* for SEZ and *Cdhr1* for M/T region); bottom: cluster labels mapped on tissue coordinates. **g,** Comparison of cell type occupancy of each spot in Ex-ST and standard Visium data (ST) visualized for two cell types, OB neuroblasts and M/T cells. **h,** Expression of olfactory receptors (Olfr) in both Ex-ST and standard Visium datasets. DAPI (Ex-ST) and HE (standard Visium) images showing the locations of glomeruli with boundaries indicated by dashed lines. The localization of Olfr, detected Ex-ST, in specific glomeruli is in agreement with coordinates provided by a previous study using serial sectioning ^23^. Scale bars for expanded samples in all panels have been rescaled by expansion ratio.

We first performed Ex-ST on the mouse olfactory bulb (MOB), in which neurons are organized in spatially distinct laminated zones with well-defined molecular signatures ^9,10^. With matched sequencing saturation (**Supplementary Table 1**), Ex-ST captured more unique molecular identifiers (UMIs) per gene from the same tissue area compared to the standard Visium protocol (**Fig. 1b**), and showed excellent reproducibility between replicates (**Extended Data Fig. 2a**). As expected, the number of unique genes detected per spot was lower using Ex-ST, but after normalizing the tissue area covering each spot, Ex-ST captured more unique genes per spot (**Fig. 1c**). These observations suggest that Ex-ST achieves higher coverage of transcripts while each spot accesses a relatively pure pool of genes.

The spatial distribution of region-specific marker genes, including *Sox11* for subependymal zone (SEZ), *Nrgn* for granular layer (GCL), *Cdhr1* for mitral (M) and tufted (T) cell layers, *Nrsn1* for glomerular layer (GL), and *Ptn* for olfactory nerve layer (ONL) ^9^, showed expected layered patterns (**Fig. 1d**, **Extended Data Fig. 2b**) ^10^, and indeed had higher UMI counts compared to standard Visium protocol (**Extended Data Fig. 2c**). By clustering the Ex-ST data, we identified 6 clusters, each corresponding to a morphological layer of MOB, annotated based on their marker genes and spatial locations (**Fig. 1e**, **Extended Data Fig. 2d-f**). In particular, SEZ and M/T clusters covered only 1-2 lines of spots, consistent with the width (< 50 μm) of these layers (**Fig. 1f**).

We next determined whether tissue expansion reduced the number of cells occupying each spot. We applied stereoscope, a deconvolution algorithm to infer cell type composition of each spot using single-cell transcriptomes ^11^. We observed that Ex-ST data contained more spots with only one dominant cell type compared to standard Visium (**Extended Data Fig. 3**). For example, SEZ and M/T regions had 16 and 96 spots, respectively, where a single cell type occupancy was over 50% with highest values reaching ~80%. In contrast, standard Visium data had no spots in SEZ and only 2 in the M/T region with dominating cell type occupancy barely over 50%, suggesting that Ex-ST provides near single-cell resolution (**Fig. 1g**).

Encouraged by the enhanced resolution of Ex-ST, we examined the glomeruli in olfactory bulbs, whose sizes range between 50 to 120 μm in diameter. Consequently, resolving gene expression within individual glomeruli using standard Visium is challenging. In contrast, Ex-ST allowed us to map olfactory receptors directly onto single glomeruli (**Fig. 1h**), consistent with previous findings that olfactory sensory neurons projecting their axons into a common glomerulus should express the same type of olfactory receptor (Olfr) ^12,13^. This result demonstrates the potential of Ex-ST to reveal tissue organization at a finer scale.

Applying Ex-ST on mouse hippocampus, we observed a similar increase in the number of captured UMIs in comparison to the standard Visium protocol (**Extended Data Fig. 4a, b**). We identified 11 clusters corresponding to different anatomical regions, expressing known marker genes (**Fig. 2a**, **Extended Data Fig. 4c-e**). Stereoscope analysis showed that most spots in the cornu ammonis (CA) and dentate gyrus (DG) regions had >80% of occupancy by a single cell type whereas the standard Visium data rarely had spots with cell type occupancy over 50% (**Fig. 2b**). Overall, Ex-ST shifted the cell type composition per spot to near single cell type occupancy throughout the hippocampus (**Fig. 2c**).

**Fig. 2:**
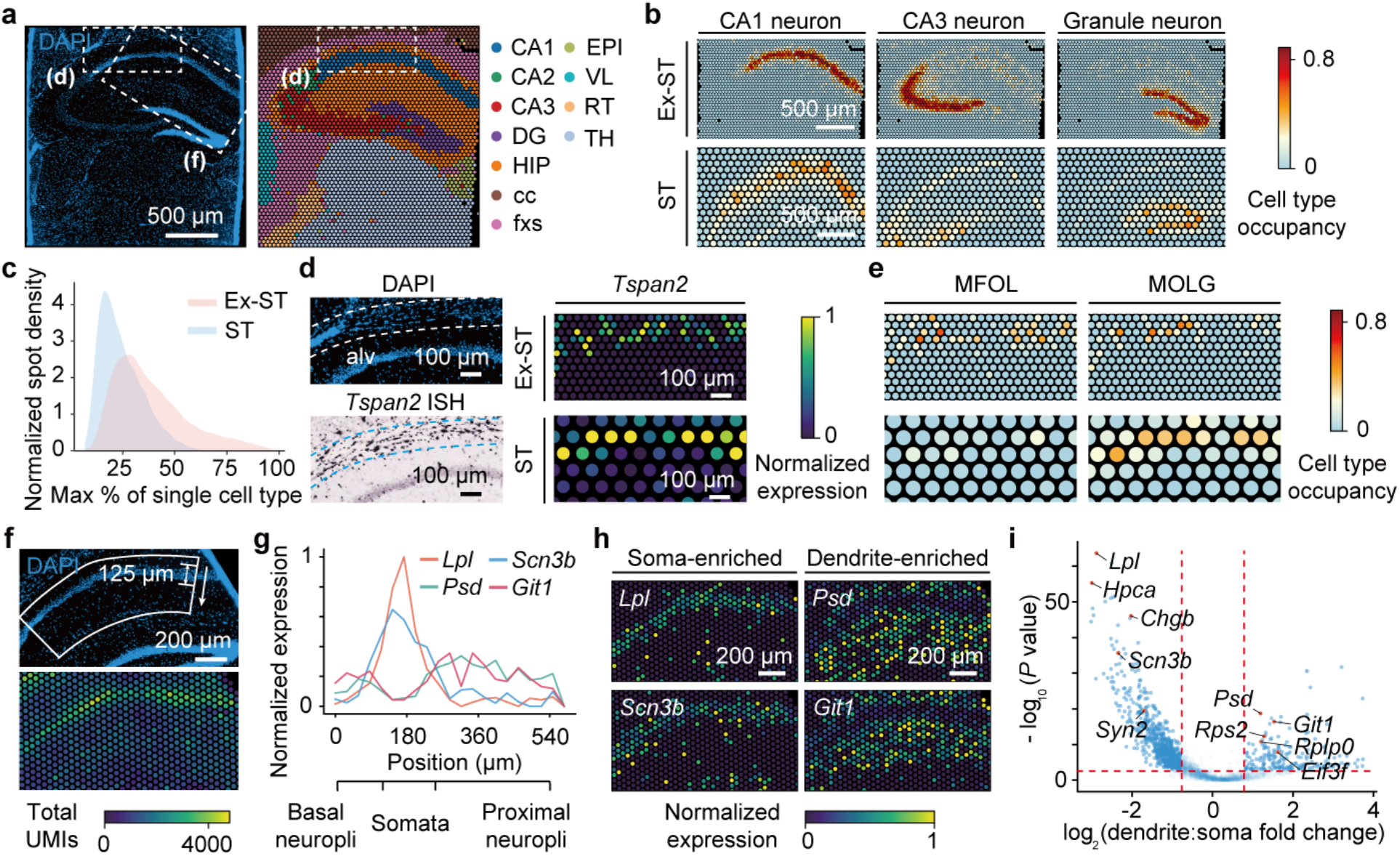
Ex-ST dissects fine structures in the mouse hippocampus. **a,** DAPI image of an expanded mouse hippocampus section (left) and annotated clusters on tissue coordinates (right): CA1-3, hippocampal subfields; DG, dentate gyrus; HIP, hippocampal region; cc, corpus callosum; fxs, fornix system; EPI, epithalamus; VL, lateral ventricle; RT, reticular nucleus of the thalamus; TH, thalamus. **b,** Comparison of cell type composition of individual spots in Ex-ST and standard Visium data visualized for three cell types. **c,** Occupancy of dominating cell type in each spot of both Ex-ST and standard Visium data visualized as a density plot. **d,** Magnified DAPI image showing the sporadic distribution of cell bodies in alveus (top left; alv, dashed outlines). ISH of *Tspan2* showing oligodendrocyte cell bodies, obtained from Allen brain atlas (bottom left) ^10^. Comparison of normalized *Tspan2* expression in Ex-ST and standard Visium data (right). **e,** Comparison of cell type composition of individual spots in Ex-ST and standard Visium data visualized for oligodendrocyte subclasses. **f,** Magnified DAPI image showing the high nuclear density within the CA1 region (top). Spatial distribution of UMIs within the CA1 region (bottom). **g,** Expression profiles of soma-dendrite differentially expressed genes along the axis indicated in **f**. **h,** Normalized spatial expression of soma-enriched and dendrite-enriched genes. **i,** Volcano plot showing differential gene expression between soma and dendritic regions. *P* values: twotailed, two-sample Mann-Whitney-Wilcoxon tests. Gene with log2 (fold change) > 0.8 and *P* value < 0.001 (dashed lines) are considered differentially expressed. Several previously known soma- and dendrite-enriched genes are specified. Scale bars for expanded samples in all panels have been rescaled by expansion ratio.

Notably, by expanding the tissue, we resolved subclasses of oligodendrocytes in alveus, a dense layer of myelinated axonal fibers covering the ventricular parts of the hippocampus. Consistent with *in situ* hybridization (ISH), oligodendrocyte markers, such as *Tspan2*, were distributed sporadically within this layer in our Ex-ST data, whereas their expression in the standard Visium was more continuous and diffusive (**Fig. 2d**, **Extended Data Fig. 5a**). Deconvolving the expression of individual spots using stereoscope revealed the distinct patterns of several oligodendrocyte subtypes in the Ex-ST data, including myelin forming oligodendrocyte (MFOL) and mature oligodendrocyte (MOLG) ^14^, which were mixed in the standard Visium data (**Fig. 2e**). This observation suggests that Ex-ST improves the capacity of distinguishing spatially-mixed subcell types that are only subtly different.

Lastly, we demonstrated that Ex-ST can measure subcellular distribution of transcripts. While most neurons have more transcripts in somata, CA1 neurons actively transport specific transcripts to dendrites ^15^. Indeed, the total number of UMIs dropped from the somata into stratum radiatum in a distance matching with the expected size of somata (~125 μm) (**Fig. 2f**), suggesting that the lateral diffusion of transcripts during mRNA capture is negligible. Meanwhile, Ex-ST separated differentially located transcripts in somata and dendrites, including dendrite-enriched genes such as *Psd* and *Git1* (**Fig. 2g, h**). Through differential gene expression analysis and after removing glial transcripts, Ex-ST identified ~500 dendrite-enriched genes that were unresolvable with the standard Visium protocol, all of which overlapped with at least one of the three previous studies using orthogonal methods to identify dendritic transcripts in CA neurons (**Fig. 2i**, **Extended Data Fig. 5b**, **Supplementary Table 2**) ^16–18^.

In summary, Ex-ST bypasses the density limitation of spatially barcoded capture arrays by expanding polyelectrolyte gels to which mRNAs are anchored. Enlarged biological structures cover more capture spots, which pushes the resolution of the Visium array from 55 to 20 μm approaching the single-cell resolution and improves RNA capture to detect more genes per area. Although we demonstrate Ex-ST using the Visium platform, this approach is orthogonal to other method development efforts in the field that have primarily focused on shrinking the capture spot size ^16,19^, thereby, can be potentially integrated with other techniques to enhance their performance as well.

## Methods

### Animals

Adult male CD1 mice (3-6 months old, Charles River) were used for the Ex-ST experiments. Animal procedures were approved by the Stanford University Animal Care and Use Committee and were in accordance with NIH guidelines. Animals were group-housed on a 12-hour light-dark cycle with food and water available *ad libitum*. Mouse olfactory bulb samples for standard Visium experiment were commercially purchased from Adlego Biomedical, which operates under ethical permission nr. 17114-2020.

### Tissue preparation

Animals were euthanized by CO_2_ followed by cervical dislocation. Brain tissue was quickly dissected and embedded in O.C.T. (Tissue tek, cat# 981385). The tissue block was snap-frozen using liquid nitrogen vapor until the O.C.T. block was hardened and stored in −80 °C until use. Tissues were cryosectioned to 15 μm thickness on a Leica cryostat and placed onto charged glass slides (VWR, cat# 75799).

### Visium slides for Ex-ST

Ex-ST used multimodal spatial arrays. Each capture area has dimensions of 6.5 × 6.5 mm, containing 5,000 spots each with a diameter of 55 um. Surface capture probes include spatial barcodes, UMIs, and 50 TVN sequence to allow for polyA tail mediated capture of mRNA molecules.

### Expansion

All steps prior to RT were performed on charged glass slides (VWR). This eliminates the uneasy task of placing a tissue section into the Visium capture area, which can result in a failed positioning or tissue folding requiring a slide reset. Sections were fixed with acetone for 1 h at -20 °C, followed by incubation in the hybridization buffer, containing 1 μM anchor probe ^20^ (a 15-nt sequence of alternating dT and thymidine-locked nucleic acid (dT+) with a 5’-acrydite modification, Integrated DNA Technologies), 2× SSC, 30% [v/v] formamide (Millipore Sigma, cat# 11814320001), 0.1% [w/v] yeast tRNA (Millipore Sigma, cat# 10109223001), 1% [v/v] RNase inhibitor (Progma, cat# N2615), and 10% [w/v] dextran sulfate (Sigma, cat# D8906), at 37 °C under saturation humidity for 40 h. After anchor probe hybridization, sections were washed three times with 1× SSC.

Expansion protocol followed a sequence of gelation, proteinase K digestion, and expansion, as previously described ^21^. Briefly, samples were washed three times in PBS, then incubated in the monomer solution (1× PBS, 2 M NaCl, 8.625% [w/w] sodium acrylate, 2.5% [w/w] acrylamide, 0.15% [w/w] N,N’-methylenebisacrylamide) for 45 min at 4°C. Gelation was initiated by adding 0.2% [w/v] ammonium persulfate (ThermoFisher Scientific, cat# 17874), 0.01% [w/w] 4-hydroxy- 2,2,6,6-tetramethylpiperidin-1-oxyl (4-hydroxy-TEMPO, Sigma-Aldrich, cat# 176141), 0.2% [w/w] tetramethylethylenediamine (ThermoFisher Scientific, cat# 17919). The gelation step was performed at 37°C for 1–2 h.

After gelation, samples were gently removed from the chamber and digested overnight at 37°C in 8 U mL^-1^ proteinase K (NEB, cat# P8107) in digestion buffer (1× TAE buffer, 0.5% Triton X-100, 0.8 M guanidine HCl). Gels were then removed from the digestion buffer and placed in 0.1× SSC to expand. 0.1× SSC was exchanged every 15 min for 3–5 times until the gel size plateaued. To stain nuclei, expanded samples were incubated with 100 μM DAPI in 0.1× SSC for 30 min. Both bright field and DAPI images of expanded samples were taken after the gel was placed on top of the Visium array and prior to mRNA release and reverse transcription (RT).

### Ex-ST library preparation

First, to determine the optimal temperature for mRNA release from the gel, several temperatures were tested. We analyzed the released mRNAs using bioanalyzer (Agilent 2100) to measure concentration and length distribution (**Extended Data Fig. 1a**). We observed that mRNAs were released from the gel with high integrity as long as temperature exceeded 41 °C (**Extended Data Fig. 1b, c**), therefore we chose to perform this step at 45 °C for 30 min to ensure complete release. RT was conducted at 42 °C for 60 min followed by 53 °C for 45 min using the Visium Spatial Gene Expression reagents (10x Genomics). This extended time of RT was chosen to enhance the reaction efficiency. After RT, the second strand synthesis was performed according to the 10x Genomics Visium Spatial Gene Expression protocol, followed by qPCR to determine the number of amplification cycles. An extra PCR cycle was added in the amplification step, in addition to the recommended number of cycles in the Visium Gene Expression protocol. A higher concentration of SPRIselect beads (0.8×) was used in the post-amplification cleanup step to retain more fragments. The rest of the library preparation was carried out according to the 10x Genomics Visium Spatial Gene Expression protocol (User Guide CG000239_RevF). Finished libraries were sequenced on a NextSeq 2000 platform (Illumina).

### Image acquisition and processing

Bright field and DAPI images of expanded samples were taken on a Zeiss Axio Observer Z1 inverted microscope equipped with an Axiocam 503M camera and a 5×/0.25 objective. The imaging time was minimized to prevent gel from drying and shrinking. Bright field image of the array frame and gel position within each capture area was aligned with the corresponding DAPI image. Manual selection of spots under the tissue was done using Loupe Browser (v 4.0.0, 10x Genomics).

### Standard Visium library preparation of MOB

The fresh-frozen MOB sample was cryo-sectioned at 10 μm thickness, placed onto Visium glass slides and stored at –80 °C prior processing. The spatial gene expression libraries were prepared according to the manufacturer’s protocol (Visium Spatial Gene Expression, User Guide CG000239_RevC). Finished libraries were sequenced on a NextSeq 2000 platform (Illumina).

### Data processing

The MOB data generated by the standard Visium protocol were processed by space ranger software (v 1.0.0, 10x Genomics). All Ex-ST data were processed by space ranger software (v 1.3.0 adapted to work with custom barcode list, 10x Genomics). Reads were aligned to the prebuilt mouse reference genome (mm10, 10x Genomics). Processed output files for the standard Visium data of the coronal mouse brain section containing the hippocampus region were downloaded from 10x Genomics website.

### Data analysis

Analysis of data generated by both Ex-ST and standard Visium protocol was performed in Python (version 3.7.12) and R (version 4.0.5). Spots with fewer than 100 genes detected (UMI > 1) were excluded from the analysis. For spatial gene expression plotting and differential gene expression analysis, expression at each spot was normalized using Scanpy (version 1.8.2) to the same total count. Dimensional reduction and clustering were performed using the Self-Assembling Manifolds (SAM) algorithm (version 0.8.9) ^22^ with default parameters. Spot deconvolution analysis was performed using stereoscope ^11^. The mouse brain single-cell RNA-seq data used for deconvolution was the “Adolescent mouse brain” dataset obtained from http://mousebrain.org/. Spatial plots for cell type mapping were produced in R (version 4.0.5), using the Seurat (version 4.1.1) and *STUtility* (version 0.1.0) R packages. CA1 soma and dendrite regions used for differential gene expression analysis were selected manually based on the DAPI images. *P*-values of gene expression differences were calculated via Mann-Whitney-Wilcoxon test using sciPy library (version 1.9.1) with default parameters. Gene expression profiles along the soma-dendrite axis were generated by averaging the expression of spots at the same distance from the soma region.

### Data and code availability

Mouse brain section containing hippocampus region is a publicly available dataset that can be found on 10x Genomics website (https://www.10xgenomics.com/resources/datasets/mouse-brain-section-coronal-1-standard-1-0-0). Single-cell RNA-seq dataset of mouse brain can be downloaded from http://mousebrain.org/adolescent/downloads.html. All data generated in this study (standard Visium data on MOB, all Ex-ST data) including spaceranger output files, stereoscope output files, DAPI and brightfield images will be available upon publication. Raw sequence data will be deposited upon publication at NCBI GEO.

## Acknowledgement

We thank Y Lim for critical discussions, and K Zhu and H Matsunami for sharing the data with us to validate the mapped glomeruli positions. YF is a Bio-X Stanford Interdisciplinary Graduate Fellow. CC is supported by a NSF Graduate Research Fellowship and a Stanford Graduate Fellowship. BW is a Beckman Young Investigator. This work is supported by an NIH grant (1R35GM138061) to BW, the Neuro-omics project of Wu Tsai Big Ideas in Neuroscience program to L Luo and BW, the European Research Council (ERC) under the European Union’s Horizon 2020 research and innovation program (grant agreement no. 101021019) to JL, and Swedish Research council to JL.

## Author contributions

YF, ZA, JL, and BW initiated and designed the project; YF, ZA, YW, and CC performed the experiments; YF and ZA analyzed the data with assistance from YW, L Larsson, and MH; YF, ZA, and BW wrote the paper with input from all other authors; L Luo, JL, and BW provided supervision and guidance.

## Extended Data Figures

**Extended Data Fig. 1:**
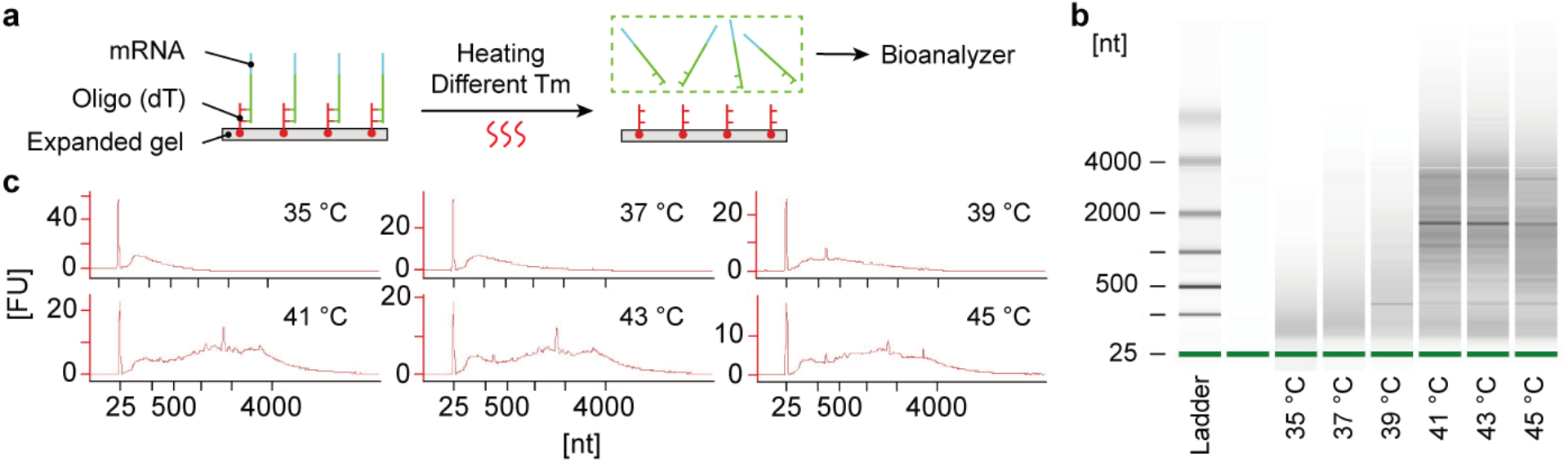
Optimization of mRNA release from the gel. **a,** The workflow to identify the optimal release temperature. Gel with anchored transcripts was heated for 30 min at different temperatures to release mRNAs, which were collected and quantified on a Bioanalyzer. **b,** Bioanalyzer gel image showing mRNAs released from the gel at different incubation temperatures. **c,** Bioanalyzer traces showing the released mRNAs. The absence of rRNA peaks is expected as these RNA should not be anchored in the gel matrix.

**Extended Data Fig. 2:**
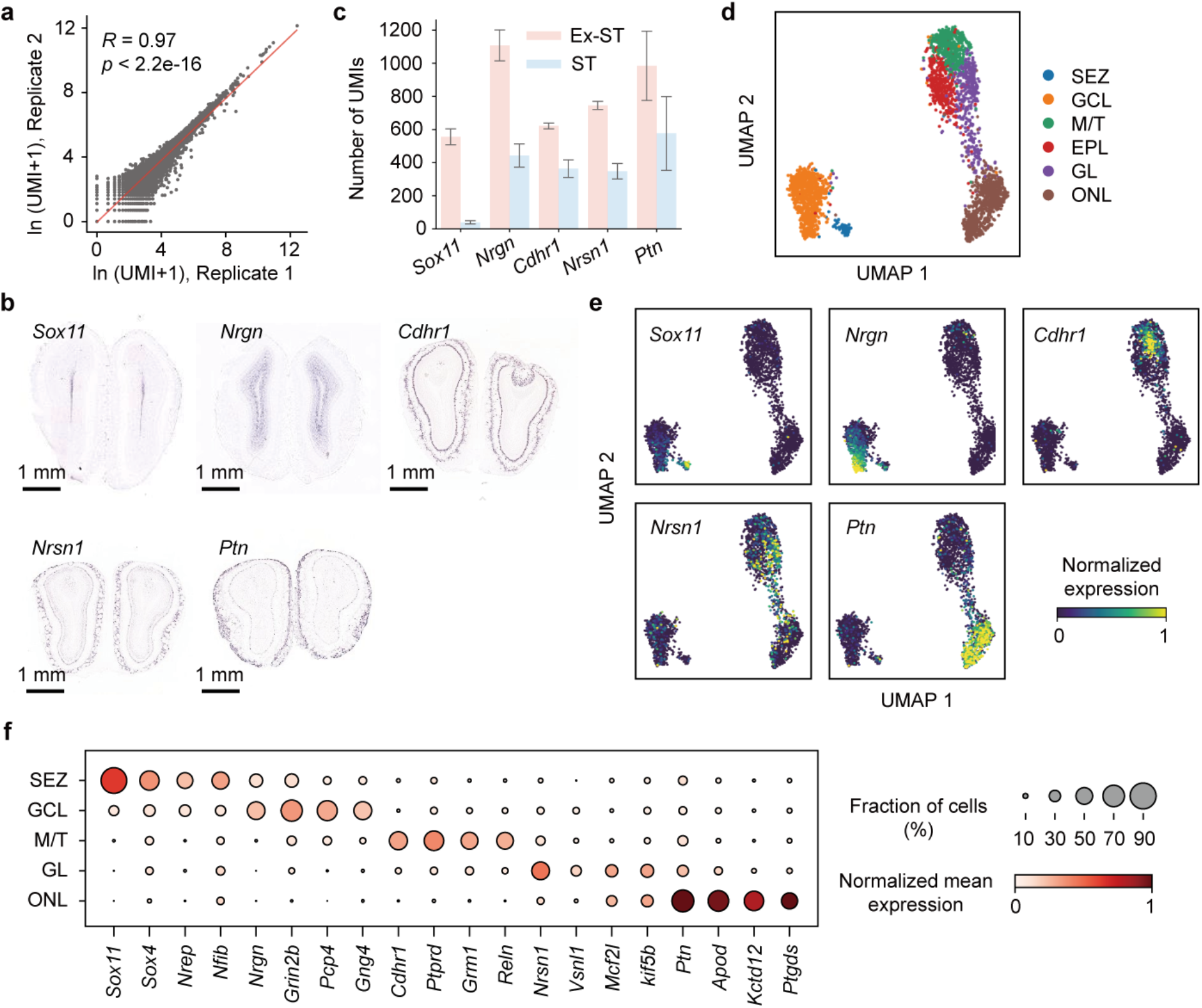
Comparison of replicates and annotation of spatial clusters of the MOB Ex-ST data. **a,** Comparison of the number of UMIs per gene between two Ex-ST replicates on MOB. **b,** The expression of region-specific marker genes detected by ISH, obtained from Allen Brain Atlas ^10^. **c,** Comparison of number of UMIs of select region-specific marker genes measured by Ex-ST and standard Visium from matched tissue areas. Error bars: range of two replicates. **d,** UMAP showing Ex-ST MOB clusters identified by unsupervised clustering, colored by cluster annotation. **e,** Marker gene expression overlaid on the UMAP visualization. **f,** Dot plot showing top marker genes for each cluster.

**Extended Data Fig. 3:**
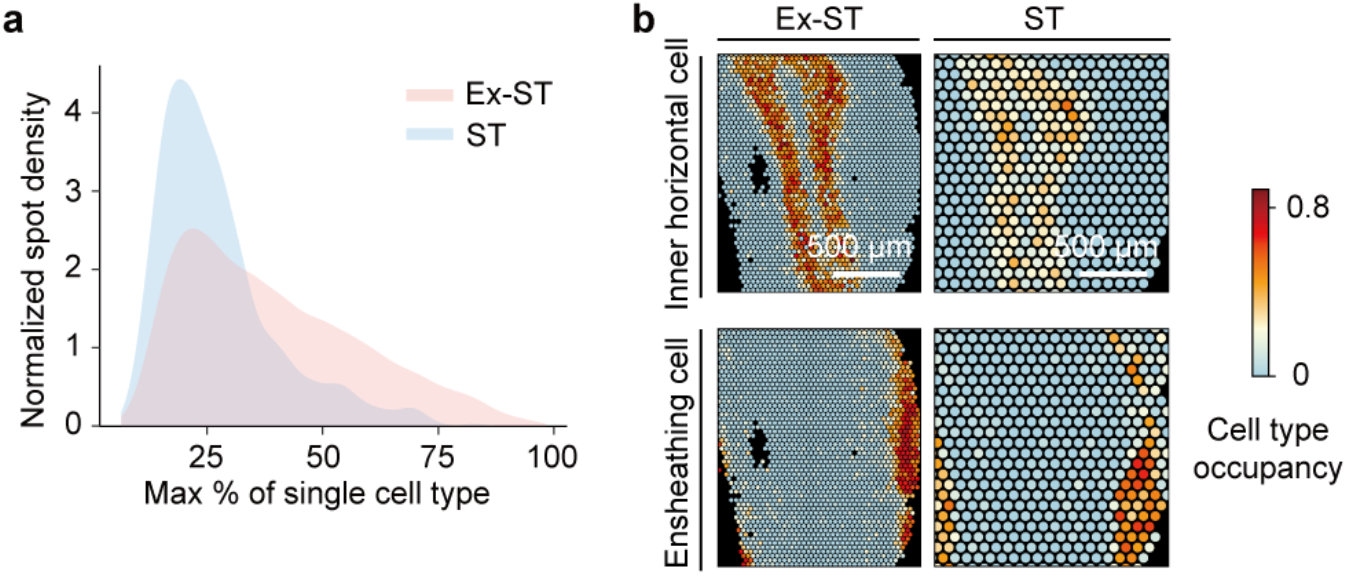
Cell type compositions of spots in the MOB datasets. **a,** Density plot showing the distribution of dominating cell type occupancy at each spot for both Ex-ST and standard Visium datasets. **b,** Comparison of cell type composition at individual spots in Ex-ST and standard Visium datasets visualized for two cell types, inner horizontal cells located in GCL and ensheathing cells in ONL.

**Extended Data Fig. 4:**
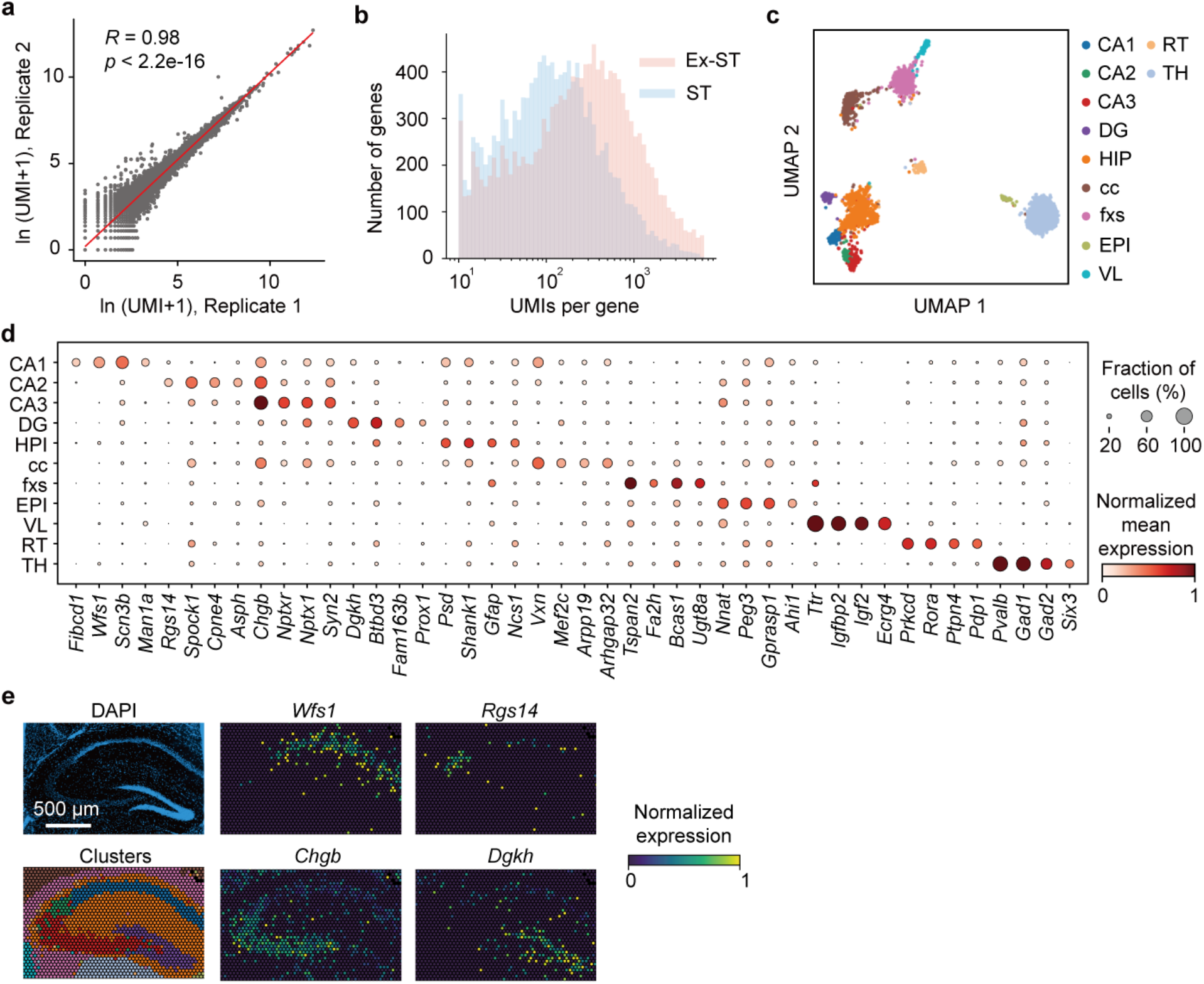
Comparison of replicates and annotation of spatial clusters of the Ex-ST mouse hippocampus data. **a,** Comparison of the number UMIs per gene between two Ex-ST replicates. **b,** Comparison of number of UMIs for all genes present in Ex-ST and standard Visium datasets, each containing one section of the hippocampal region. **c,** UMAP showing annotated clusters identified by unsupervised clustering of Ex-ST hippocampus data. **d,** Dot plot of top marker genes for each cluster. **e,** Spatial gene expression of region-specific marker genes measured by Ex-ST, i.e., *Wfs* for CA1, *Rgs14* for CA2, *Chgb* for CA3, *Dgkh* for DG.

**Extended Data Fig. 5:**
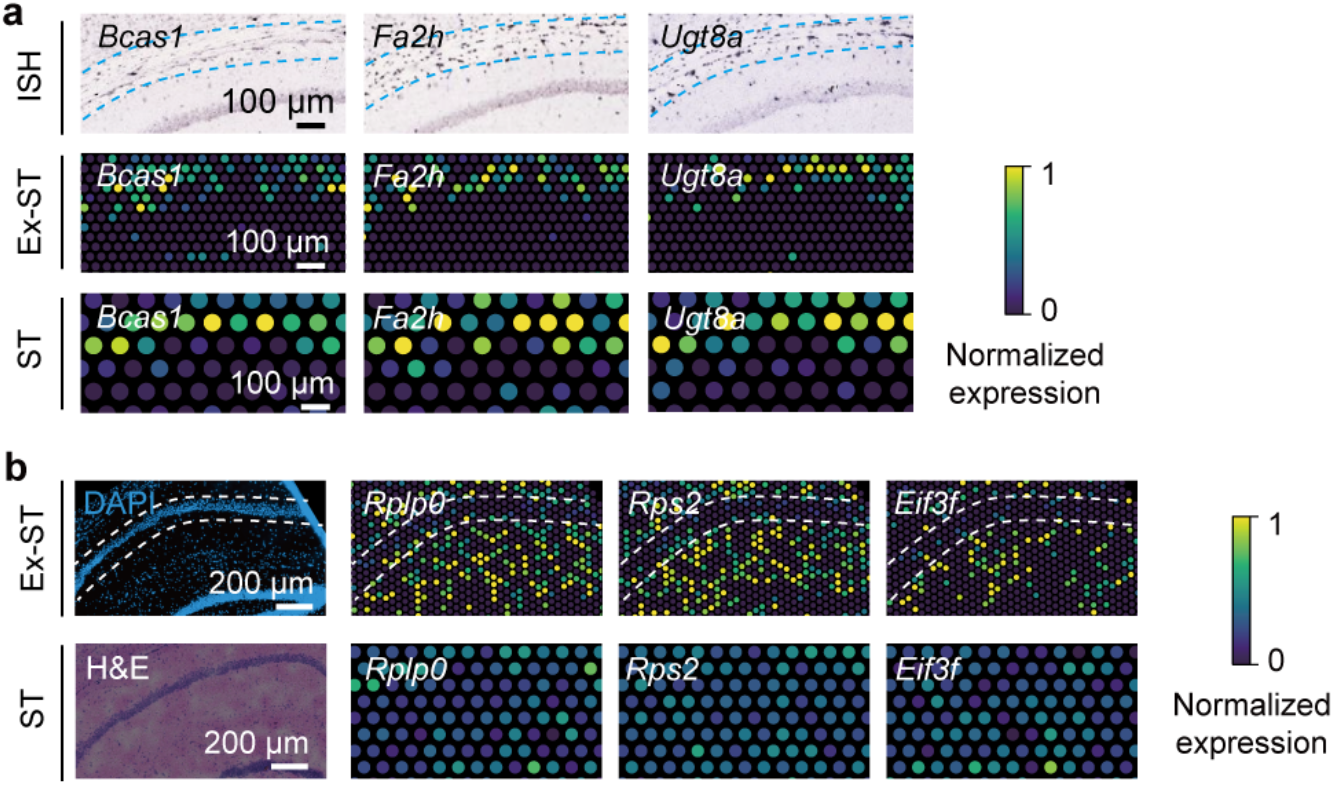
Ex-ST detects oligodendrocyte marker genes and dendrite-enriched transcripts. **a,** Comparison of ISH (top), obtained from Allen Brain Atlas ^10^, normalized spatial expression of oligodendrocyte marker genes detected by Ex-ST (middle) and standard Visium (bottom). **b,** Comparison of normalized spatial distribution of lowly-expressed dendrite-enriched transcripts between Ex-ST and standard Visium. Note the lack of pattern in the standard Visium data.

**Supplementary table 1.**
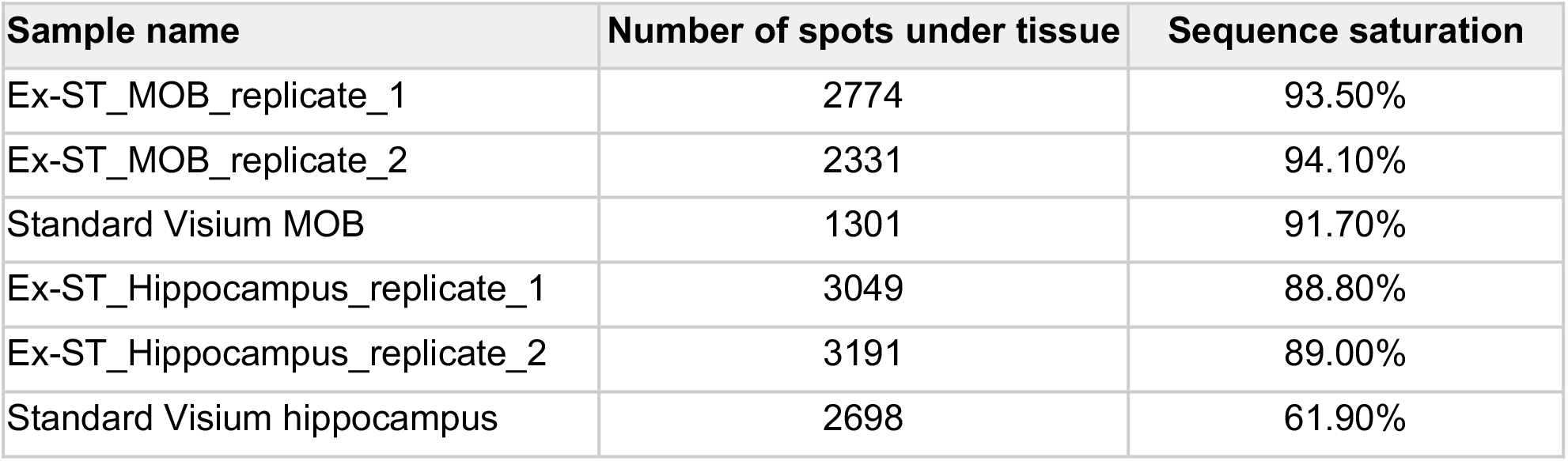
Sample information obtained from space ranger software (10x Genomics). Standard Visium MOB dataset is collected on a tissue section containing two bulbs. Both bulbs are combined to compare with expanded MOB data with two replicates each containing one bulb. Hippocampus data includes matched tissue areas from coronal sections of the mouse brain in the hippocampal region.

**Supplementary table 2.** Dendrite-enriched transcripts detected by Ex-ST but unresolvable using the standard Visium protocol. Also provided are the previous measurements ^16–18^ that support the dendritic expression of these transcripts.

